# DextraDemixer enables accurate identification of antigen-specific T cells from pMHC multimer experiments

**DOI:** 10.64898/2026.06.23.733339

**Authors:** Yang An, Felix Drost, Irene Bonafonte-Pardàs, Myriam Grotz, Kilian Schober, Benjamin Schubert

## Abstract

Antigen specificity of T cells defines the adaptive immune response, yet the vast majority of known T cell receptors (TCRs) lack annotated antigen targets. Single-cell peptide-MHC (pMHC) multimer assays offer a scalable approach to map TCR-antigen interactions. Still, their utility is limited by pervasive non-specific binding and severe overlap between signal and noise, which confound the accurate identification of antigen-specific cells. To address these limitations, we present DextraDemixer, a Bayesian hierarchical mixture model that disentangles antigen-specific T cells from background noise in pMHC multimer data. The model integrates information from negative controls and clonotype structure while providing calibrated uncertainty estimates for classification. We further introduce a dynamic thresholding scheme that enables credible interval-bounded control of the false discovery rate. Extensive benchmarking on simulated datasets and antigen-specific spike-in experiments demonstrated the model’s robustness and improved accuracy over established methods. In a longitudinal SARS-CoV-2 vaccine study, DextraDemixer identified antigen-specific TCRs characterized by high sequence similarity, elevated antigen-specificity prediction scores, and strong clonal purity. Annotations showed high concordance with external validation data and supported the identification of antigen-specific motifs. Overall, DextraDemixer provides a principled probabilistic framework for reliable identification of antigen-specific TCRs from single-cell pMHC-multimer assays.

## Introduction

Identifying the antigen specificity of T cells is fundamental for understanding the adaptive immune system’s behavior in disease settings and its contribution to homeostasis. The specificity of a T cell is determined by the variable domain of its heterodimeric antigen-recognizing T-cell receptor (TCR), particularly the complementary determining regions (CDR1-3). These CDRs interact with antigenic peptides, called epitopes, and the major histocompatibility complex (MHC) presenting the epitope, forming the TCR-pMHC complex^1^. Recent advances in single-cell sequencing allow large-scale identification of antigen-specific T cells through pMHC-multimer assays, combined with a detailed map of the cell’s transcriptional state and full-length TCR sequence^2–9^. In these assays, pMHC multimers carrying unique oligonucleotide barcodes label T cells whose receptors bind a given pMHC complex, allowing antigen specificity to be linked directly to cellular phenotype and TCR sequence. These combined assays have become an important technology in T-cell immunology, enabling the discovery and detailed characterization of antigen-specific T-cell populations in vaccination^10–13^, infection^14,15^, and in cancer^16–18^, among others.

However, these high-throughput assays suffer from low signal-to-noise ratios, non-specific binding, the occurrence of contamination, and sequencing dropout, all of which confound reliable identification of antigen-specific T cells. As a result, most common data processing approaches require a high degree of manual analysis to find appropriate pMHC-UMI thresholds, along with additional experimental verification on a subset of potential epitope-specific T cells^11^. Beyond thresholding absolute UMI counts^11,19^, Minervina et al. employed relative thresholds, normalized across all pMHCs^14^. Similarly, Francis et al. applied inter-multimer and -cell z-score normalization to correct the pMHC signal and classified cells with MHC-specific thresholds calibrated using additional experimental data^20^. Zhang et al. introduced ICON^21^, an analysis pipeline that combines manual intervention with automated quality-control filters, noise correction using negative control multimers, clone-aware signal correction, cell- and multimer-wise normalization of corrected UMIs, and empirically determined specificity assignment thresholds. Similarly, the ITRAP pipeline^22^ by Povlsen et al. applies cascaded filtering steps, including auxiliary information like hashing singlets, and employs clonotype-pooled analyses to automatically optimize UMI thresholds, balancing high concordance between pMHC assignments and clonotype-based specificity with minimal loss of unassigned cells. The most recent method, the BEAM^23^ algorithm developed by 10x Genomics, employs a different strategy by utilizing a fully probabilistic model. It applies a Beta distribution parameterized by the selected pMHC and negative control UMI counts as a proxy for the noise distribution to compute a continuous antigen-specificity score that requires subsequent manual thresholding.

Despite these advances, existing methods either rely on arbitrarily chosen parameters, do not exploit clonotype identity to inform antigen assignment, or fail to capture the distributional shift between negative control and noise UMI counts often observed in such experiments. To address these shortcomings, we developed DextraDemixer (**Fig. 1**), a Bayesian hierarchical negative binomial mixture model that can jointly model measured pMHC-UMI counts with negative controls, aggregates cell-specific antigen assignment probabilities across clonotypes, and provides posterior probabilities as quantitative measures of assignment uncertainty. It further supports expected FDR control through an automated thresholding procedure, reducing the need for dataset-specific manual interventions such as visual inspection of UMI distributions, empirical threshold selection, and ad hoc filtering. Together, these features make DextraDemixer a tool for automated, robust, and reproducible identification of antigen-specific T cells from pMHC-multimer assays.

**Figure 1.**
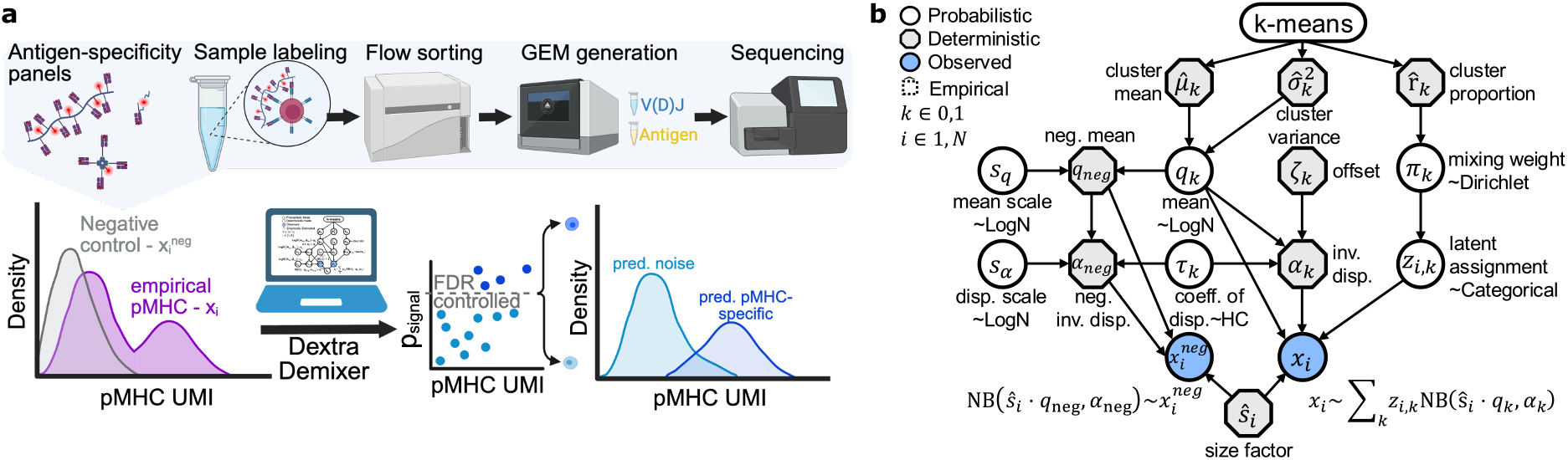
DextraDemixer pipeline. **a**, Schematic illustration of a typical pMHC-multimer sequencing workflow, resulting in the UMI count distribution for the pMHC, and potentially negative control UMI counts and TCR sequences. Due to biological and technical noise, the binder status of T cells cannot be unambiguously determined. DextraDemixer uses the pMHC UMI count, and optionally the cells’ clonotype and negative control UMI count, to infer a binder probability for each T cell to disentangle specific binders from noise. **b**, Graphical model of DextraDemixer. Abbreviations: inv., inverse; disp., dispersion; neg., negative control; LogN, log normal; NB, negative binomial; coeff., coefficient. Created with BioRender.com.

## Results

### DextraDemixer is a principled Bayesian framework for identifying antigen-specific T cells

DextraDemixer is a family of hierarchical Bayesian models that offers a principled approach to robustly isolate true binding signals from technical and biological noise in pMHC-multimer assays (**Methods - Model description; Fig. 1a, b**). For each pMHC multimer, DextraDemixer assumes that the measured UMI counts reflect two overlapping populations of cells: cells with non-specific background binding and cells with true antigen-specific binding. These two populations are modeled with separate negative binomial distributions. After inferring the parameters of both distributions from the data, the model assigns each cell a posterior probability of belonging to the antigen-specific signal component. To guide the model towards plausible background and signal components and stabilize inference in the mixture model, we derive data-driven priors using a k-means clustering step as an empirically grounded initialization. Additionally, jointly modeling observed negative control data allows calibration to the assay-specific noise profile. Crucially, rather than treating the negative control as a precise proxy of background noise, we account for probabilistic shifts in both mean and variance. By propagating uncertainty throughout the hierarchy, the model computes a statistically rigorous posterior probability of binding for each cell. Finally, leveraging the immunological premise that T cells of the same clonotype recognize the same antigen, DextraDemixer aggregates cell-level probabilities to output a clone-level median. This integration of statistical modeling with biological domain knowledge provides a highly robust probabilistic assignment, ensuring confident identification of antigen-specific populations.

### DextraDemixer robustly identifies antigen-specific cells across a wide range of scenarios

To systematically evaluate DextraDemixer, perform ablation studies, and its robustness across diverse scenarios, we developed a parameterized simulation framework (**Methods - Simulation; Supp. Fig. 1**). The framework generates UMI counts independently for each pMHC mirroring the empirical distributions, noise profiles, and clonotype structures observed on real human PBMC and murine spleen single-cell pMHC-multimer data from 10x Genomic^19^ (**Methods - Simulation; Supp. Fig. 2**). We generated simulated datasets spanning a comprehensive parameter grid comprising total cell counts (100-10,000), fraction of binders (0.001-0.5), and signal-to-noise ratios (10-200, defined as ratio of component means). These parameter ranges over multiple magnitudes captured both average cases, as well as edge cases exhibiting severe overlap between signal and noise distributions (**Supp. Fig. 3**).

We compared DextraDemixer and its extensions against BEAM, the only existing method designed to predict antigen-specific binders in a single pMHC-multimer setting. Across this comprehensive benchmark with 2,280 simulated datasets, the base version of DextraDemixer significantly outperformed the established method BEAM in all classification metrics: F1 score, Precision, Recall, and Average Precision Score (APS) (**Fig. 2a; Supp. Table 1**). The average F1 score increased from 0.67 for BEAM (95% Confidence Interval (CI): 0.65-0.68) to 0.86 for DextraDemixer (95% CI: 0.85 - 0.87; paired, two-sided Wilcoxon signed-rank test, p-value < 10^-236^). Although recall was comparable between BEAM and DextraDemixer, precision showed a large absolute increase of 24.9% for DextraDemixer. This indicates that DextraDemixer found a similar proportion of true antigen-specific cells, while producing fewer false-positive annotations, hence, a cleaner set of antigen-specific T cells. BEAM leverages the ratio between the target pMHC and negative control UMI counts for each cell to calculate its antigen-specificity score. Because measured negative controls frequently show poor alignment with the endogenous noise distribution, as observed in real datasets (**Supp. Fig. 2a**), this assumption yields error-prone classifications. By explicitly modeling this distributional shift, DextraDemixer’s negative control integration further improved its classification performance, increasing the average F1 score from 0.86 (95% CI: 0.85 - 0.87) for the base model to 0.87 (95% CI: 0.86 - 0.88; p-value < 10^-27^) (“+neg.” in **Fig. 2a**). Extending the prediction from cell-level probabilities to clonotype median aggregated probabilities provided an additional significant improvement, increasing the average F1 score to 0.90 (95% CI: 0.89 - 0.91; p-value < 10^-126^) (“+clone” in **Fig. 2a**). Ultimately, the full DextraDemixer model incorporating both negative control and clonotype-level median aggregation achieved the highest overall accuracy, with an average F1 score of 0.91 (95% CI: 0.90 - 0.92; p-value < 10^-21^).

**Figure 2.**
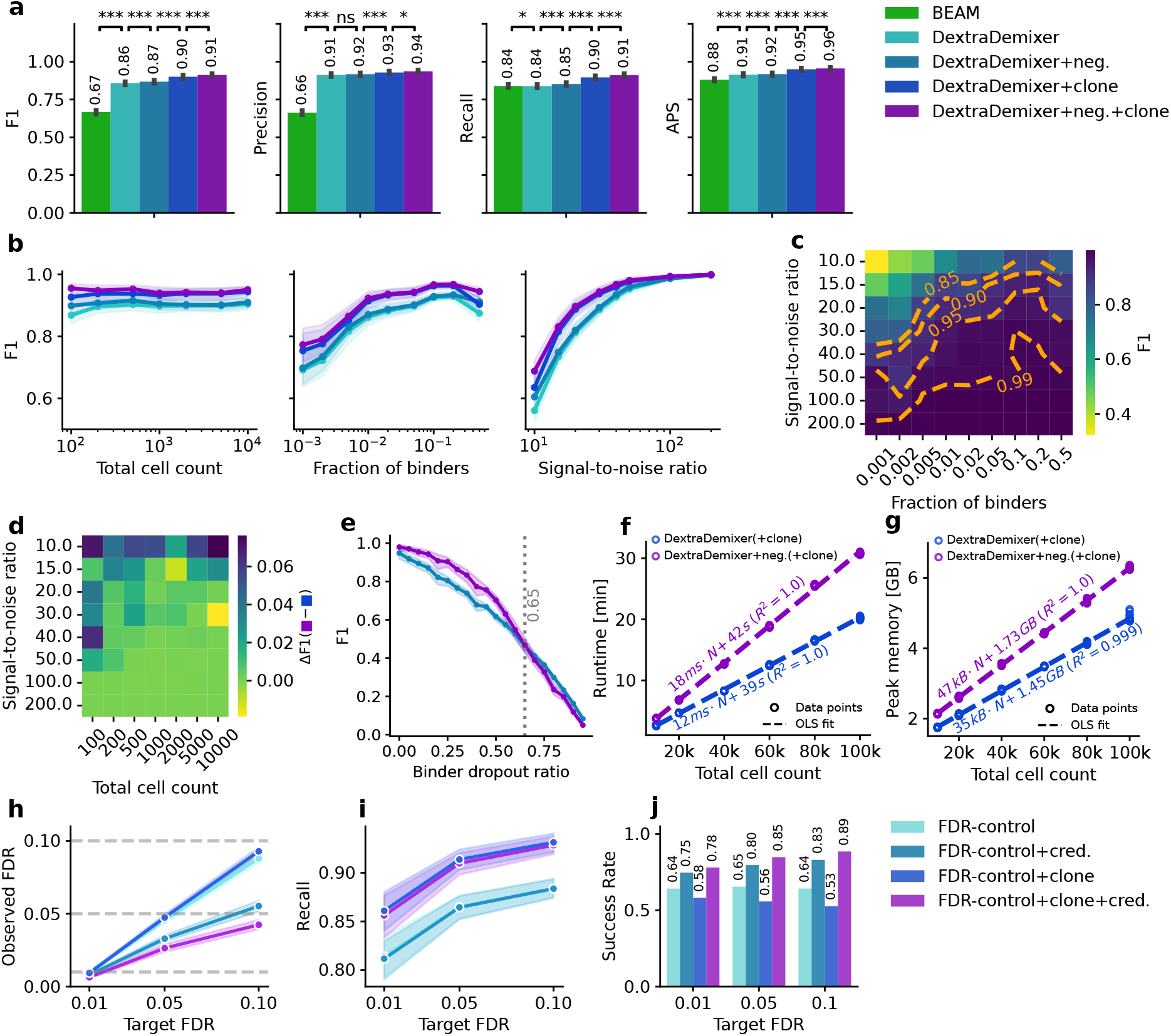
Performance on synthetic benchmark. **a**, Average F1 score, Precision, Recall, and average precision score (APS) between different variants of DextraDemixer compared to BEAM on the synthetic benchmark with *n = 2,280* datasets. Statistical significance was tested by a paired, two-sided Wilcoxon signed-rank test (ns: not significant, *: *p<0*.*05*, **: *p<0*.*01*, ***: *p<0*.*001*). **b**, Performance indicated by F1 score of DextraDemixer variants on varying simulation parameters: total cell count, fraction of binders, and signal-to-noise ratio (ratio of component means). **c**, Average F1 scores of the full *DextraDemixer+neg*.*+clone* model on a grid with different fractions of binders and signal-to-noise ratio combinations. The orange dashed lines indicate the iso-contours where the model achieves the specified F1 score on simulation parameter combinations. **d**, Difference in F1 score between the clone model with and without the negative control joint modeling. **e**, Relationship between binder dropout ratio (fraction of true binder cells exhibiting noise pMHC-UMI measurement) and classification performance with and without clonotype median-aggregated scoring. Grey line indicating the intersection between both model performances at a binder dropout ratio of 0.65. **f, g**, Dashed lines depict the ordinary least squares (OLS) fitted regression on the computational complexity with growing dataset size for **f**, runtime, and **g**, peak memory. **h, i, j**, DextraDemixer with automatic thresholding for false discovery rate (FDR)-control showing **h**, Target FDR vs. observed FDR, **i**, Target FDR vs. Recall, and **j**, Target FDR vs. Success Rate, defined as the ratio of datasets with observed FDR below the target FDR. Error bars and shaded regions depict the 95% confidence interval for bar and line plots, respectively. *+neg*.: joint negative control modeling, *+clone:* clone median aggregation. *+cred*.: credible interval-bounded FDR-control.

To systematically analyze the model’s robustness, we stratified the performance by simulation parameters (**Fig. 2b**). To isolate the effect of the number of cells while retaining the same sets of values for the other simulation parameters, datasets with a fraction of binders smaller than 0.01 were excluded, as this would yield trivial zero binders at the minimum dataset size of 100 cells. The performance remained stable across all generated dataset sizes, with the largest difference in average F1 score between any two dataset sizes within the same model being 0.04. We observed a pronounced impact of the fraction of binder parameter, with an increase in F1 score with a higher fraction of binders and a maximum difference of 0.24. However, the most dominant effect was driven by the signal-to-noise ratio, with an average F1 score of 0.56 at an extremely low signal-to-noise ratio of 10 to an F1 score of 1.0 at the highest ratio. This is expected, as severe overlaps between signal and noise distributions occur at the lower ratios and diminish with increasing values (**Supp. Fig. 3**). Analyzing interaction effects between the fraction of binders and signal-to-noise ratio revealed that performance losses induced by the low fraction of binders were mitigated by a high signal-to-noise ratio (**Fig. 2c**). While the overall improvement of the negative control extension was modest, it was particularly pronounced in challenging low signal-to-noise ratios and small dataset size regimes (**Fig. 2d**).

To test the robustness of the clonotype extension against biological dropout, we simulated scenarios where cells within binding clones randomly fail to exhibit antigen-specific binding patterns. Performance generally dropped with increasing levels of dropout. The clonotype extension tolerated up to 65% of outliers before degrading below the base model’s performance (**Fig. 2e**).

Lastly, to confirm the computational scalability of DextraDemixer for large-scale datasets, we captured the time and memory requirements on a single Intel Xeon Gold 6142M core with a dataset size grid between 10,000 and 100,000 cells. Both metrics scaled linearly with the number of cells (**Fig. 2f, g**). Notably, joint modeling of negative control UMI counts increased the per-cell runtime slope from 12 ms/cell to 18 ms/cell and the memory footprint slope from 35 kb/cell to 47 kb/cell. Processing of 100,000 cells with joint modeling of negative controls. required a peak RAM footprint of only 6.4 GB while taking 30.8 min, making the method feasible to run locally on average consumer laptops.

### DextraDemixer reliably controls the expected false discovery rate

We next sought to develop an automated thresholding strategy that controls a user-defined false discovery rate (FDR). Although a posterior probability threshold of 0.5 provides a natural classification boundary that balances Type I and II errors, downstream analyses of antigen-specific T cells are often disproportionately affected by false positives. DextraDemixer addresses this by leveraging its calibrated posterior probabilities to employ a dynamic assignment strategy to control the FDR. Thus, rather than relying on a fixed probability cutoff, DextraDemixer dynamically adjusts the decision boundary to control the expected FDR at a user-defined level (**Methods - Model**).

We validated DextraDemixer’s FDR control on a benchmark subset, constrained to samples containing at least 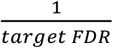 binders, ensuring a single false positive would not immediately violate the target FDR. DextraDemixer successfully constrained the average empirical FDR below the target FDRs of 0.01, 0.05, and 0.1 (**Fig. 2h**). Expectedly, relaxing the target FDR increased the recall from 0.86 at a target FDR of 0.01, up to 0.93 at a target FDR of 0.1 with the clone model (**Fig. 2i**). Moreover, leveraging DextraDemixer’s full posterior distribution for credible interval-bounded FDR control further lowered the corresponding empirical FDR and increased the success rate, while maintaining a similar level of recall (“+cred.” in **Fig. 2h-j**).

### DextraDemixer accurately identifies rare epitope-specific cells in spike-in experiments

Having established DextraDemixer’s performance in simulations, we next evaluated its ability to resolve antigen specificity in a multi-pMHC setting using empirical single-cell sequencing data with known ground truth specificity annotations using the clonal spike-in experiment reported by Gemünd et al.^25^. In this study, three spike-in T cell clone sets with defined pMHC specificities were added to HLA-mismatched peripheral blood samples: NLVPMVATV:HLA-A*02:01-specific clones recognizing the DexA dextramer, CRVLCCYVL:HLA-C*07:02-specific clones recognizing DexC, and TPRVTGGGAM:HLA-B*07:02-enriched clones predominantly detecting DexB. The three clone sets were mixed at a 1:1:1 ratio and jointly spiked in at total frequencies of either 5% or 50% of all T cells. BD Rhapsody-based single-cell immune profiling combined with pMHC-multimer sequencing was performed for 10,000 loaded T cells using the three spike-in-specific dextramers and a fourth dextramer carrying a negative control epitope that was HLA mis-matched to the spike-in clones (**Fig. 3a**).

**Figure 3.**
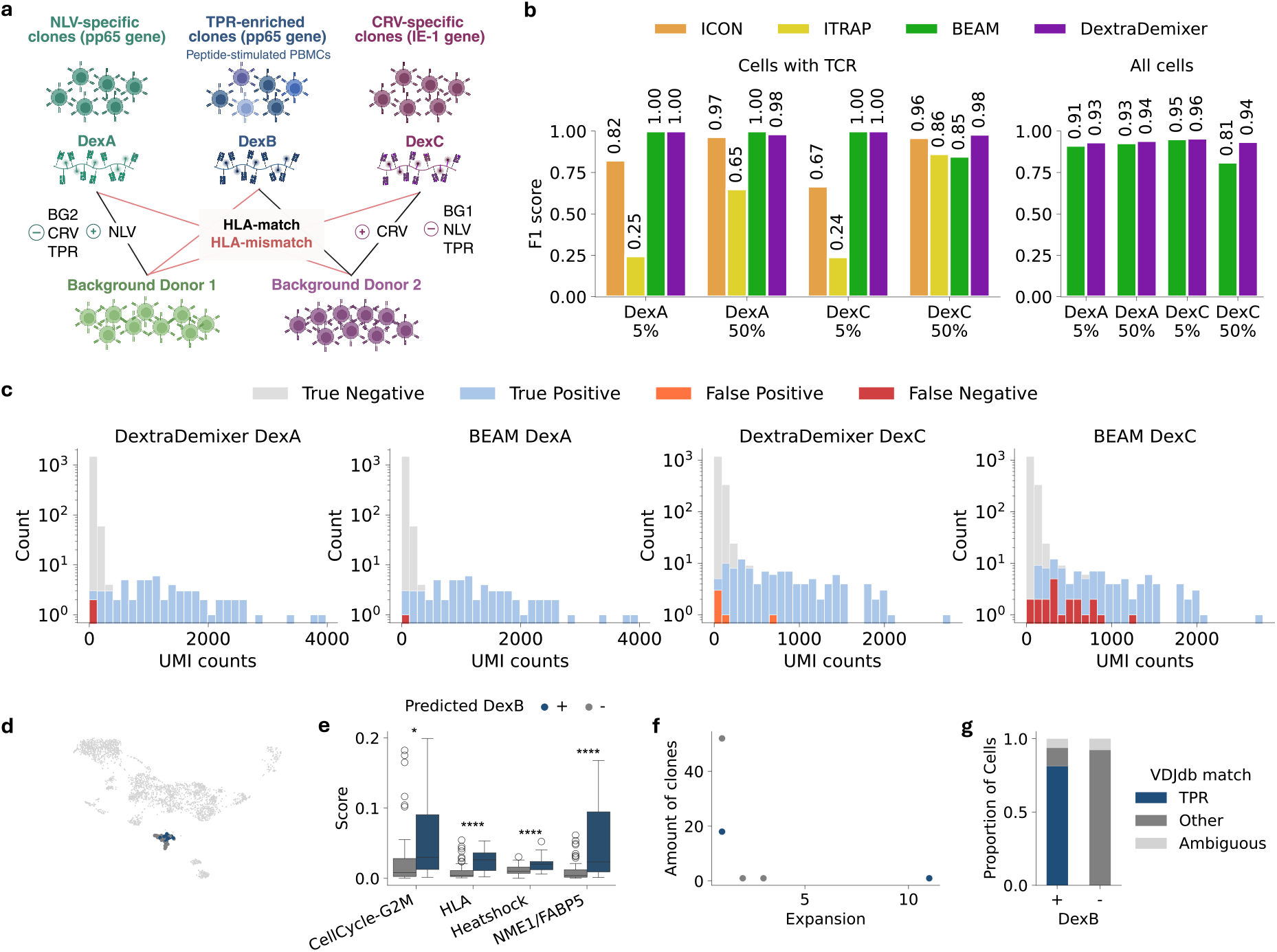
Performance on spike-in experiment dataset. **a**, Experimental design. Two spike-in clone sets with known specificity for NLV and CRV, respectively, and a TPR enriched set, were mixed into background T cells from two donors at combined spike-in frequencies of 5% and 50%. HLA-mismatched background and spike-in T cells were used as negative labels for each corresponding spike-in clone set. **b**, Comparison of DextraDemixer with negative control and clonotype extension performance to other methods across the two dextramers with known specificity and spike-in frequency settings, for cells with TCR information and all cells. For the all-cell analysis that do not require clonotype aggregation are shown. **c**, Count distribution colored by classification results for cells with TCR. **d-g**, Comparison of CD8^+^ TPR enriched cells classified as specifically and non-specifically binding DexB by DextraDemixer. **d**, UMAP colored by classification results. **e**, TCAT^24^ estimated level of antigen-activation in DexB^+^ cells compared to DexB^-^. * indicates Benjamini-Hochberg FDR-corrected p-values from a two-sided T-test. *: *FDR < 0*.*05*, ****: *FDR < 10*^*-5*^. **f**, Clonal expansion in DexB^+^ and DexB^-^ cells. **g**, Specificity of DexB^+^ and DexB^-^ cells based on VDJdb matches. Only cells with a match are included. “Other” represents cells matching any epitope other than TPR and “ambiguous” cells matching TPR and any other epitope. Created with BioRender.com

For benchmarking, we analyzed four subsets corresponding to DexA 5%, DexA 50%, DexC 5%, and DexC 50%. In each subset, the DexA- and DexC-specific spike-in cells were treated as true positives, respectively, whereas HLA-mismatched background cells and spike-in cells with alternative specificities were treated as true negatives. We benchmarked the full DextraDemixer model, including negative-control and clonotype integration, against BEAM^23^, ICON^21^, and ITRAP^22^. Because ICON and ITRAP both require complete TCR information and a multi-pMHC multimer setup, they could only be evaluated in this benchmark and only after filtering for cells with fully annotated TCRs. After filtering, the datasets contained 29 specific cells out of 587 cells in DexA 50% setting, 7 out of 835 in DexA 5%, 49 out of 703 DexC-specific cells in DexC 50%, and 3 out of 709 in DexC 5%.

DextraDemixer achieved the highest average performance with F1 of 0.99 (95% CI: 0.97-1.00), with near-perfect classification across all settings (F1≥0.98, **Fig 3b, Supp. Table 2, Supp. Fig. 4a**). BEAM reached an average F1 of 0.96 (95% CI: 0.84-1.00) with perfect predictions in three out of four settings, but was less performant in the DexC 50% spike-in setting, where antigen-specific and non-specific signal distributions overlapped most strongly (F1=0.85, **Fig 3b,c**). ICON performed well in this setting (F1=0.96) however only reached a mean F1 of 0.86 (95% CI: 0.63-1.00). Because ICON relies heavily on clonotype-level aggregation, it underperformed in the 5% spike-in settings, where positive cells were largely derived from unexpanded clones (F1=0.82 for DexA and 0.62 for DexC, **Supp. Fig. 4b**). ITRAP, also relying on clonotype aggregation, showed the lowest average performance with F1 of 0.50 (95% CI: 0.00-0.99) and, like ICON, suffered particularly in the low-data regimes (F1=0.25 for DexA and 0.24 for DexC).

To assess robustness to increased noise, we next applied the full DextraDemixer model and BEAM to all cells, including those with orphaned or unsequenced TCRs (**Fig. 3b**). Across DexA 50%, DexA 5%, and DexC 5%, both models performed similarly, with a small advantage for DextraDemixer with an average F1 of 0.94 (95% CI: 0.92-0.97) vs. 0.93 (95% CI: 0.88-0.98) for BEAM. In the more challenging DexC 50% setting, however, BEAM again showed a marked performance drop, whereas DextraDemixer remained robust (F1=0.81 vs. 0.94).

Finally, we asked whether DextraDemixer could support downstream biological interpretation by applying our model to TPR-enriched CD8^+^ cells, which consisted of a mix of specific and non-specific cells to DexB. DextraDemixer identified 29 DexB^+^ and 57 DexB^-^ cells (**Fig. 3d**). Compared to DexB^-^ cells, antigen-specific cells significantly upregulated T-cell gene programs associated with TCR-dependent activation^24^, consistent with productive antigen recognition rather than non-specific binding (**Fig. 3e**). In addition, whereas no DexB^-^ clonotype contained more than three cells, DexB^+^ cells included an expanded clonotype previously reported to recognize the TPR-peptide^25^, further supporting that DextraDemixer recovers functionally active antigen-specific clones rather than isolated false-positive (**Fig. 3f**). Orthogonal validation against VDJdb^26^ provided additional support for DextraDemixer’s annotation: 81% of DexB^+^ cells matched known TPR-reactive TCRs, with no matches in DexB^-^ cells (**Fig. 3g**).

Taken together, these results demonstrate that DextraDemixer can robustly identify antigen-specific cells across diverse experimental settings, including low signal-to-noise conditions and extremely sparse antigen-specific populations in which other methods underperformed in at least one scenario. In DexC 5% containing only three positive cells, DextraDemixer was able to recover all, demonstrating the reliable identification of rare antigen-specific cells. Beyond accurate classification, DextraDemixer recovered coherent transcriptional activation programs and clonal expansion patterns, supporting its ability to distinguish true antigen-specific T cells from non-specific pMHC-multimer signals.

### Application to SARS-CoV-2 vaccination experiments reveals coherent antigen-specific TCR signatures

Finally, we applied DextraDemixer to analyse the longitudinal single-cell SARS-CoV-2 vaccine dataset from Kocher et al.^11^. The dataset comprises 53,907 cells from 14 donors across three experiments after vaccination and utilized the 10x Genomics platform for paired sequencing of gene expression and TCR. The cells of each donor were stained with multiplexed pMHC dextramers loaded with corresponding HLA-matched SARS-CoV-2 epitopes (**Fig. 4a**). Because the dextramer panel targeted multiple epitopes and the cells originated from an enrichment rather than a purification strategy, only a small fraction of cells was expected to be specific for any individual epitope.

**Figure 4.**
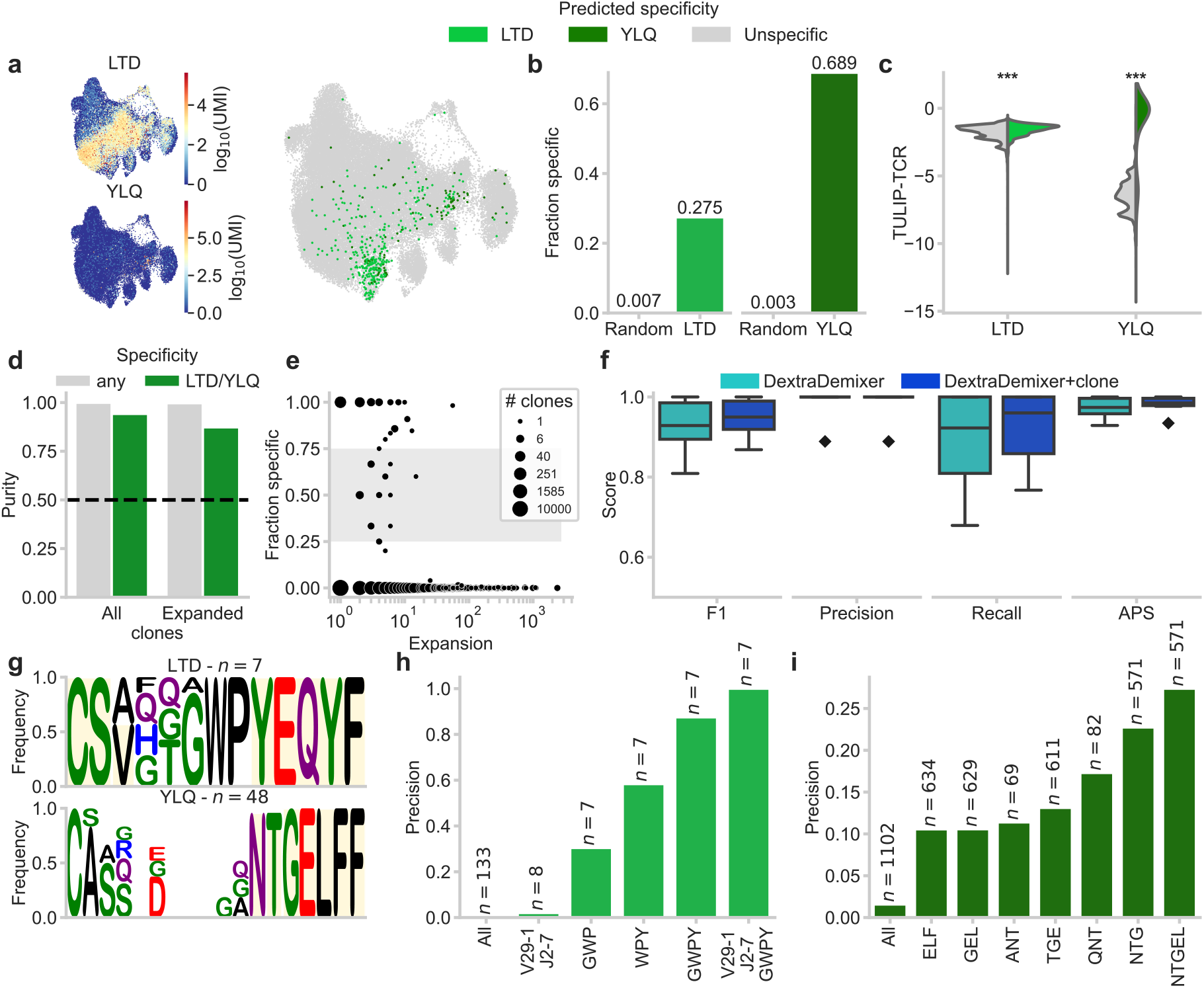
Application of DextraDemixer on SARS-CoV-2 single-cell experiments. **a**, Left, UMAP visualisation of log-transformed UMI counts for LTD and YLQ (cells without staining in gray). Right, the annotation by DextraDemixer’s cell-level variant. **b**, Fraction of antigen-specific clones for which the nearest sequence neighbor is in the same predicted specificity compared to their fraction in the dataset as a random baseline. **c**, TULIP-TCR prediction scores on DextraDemixer annotated antigen-specific and other cells. Statistical significance was tested by an unpaired, two-sided *t*-test (***: *p<0*.*001*). **d**, Average clonal purity of all clones, expanded clones, clones with at least one antigen-specific cell, and expanded clones with at least one antigen-specific cell. **e**, Relationship between expansion and fraction of antigen-specific annotation of clones. An interval of ambiguous annotation is highlighted in gray between 25% and 75%. **f**, Classification metrics based on external labels comparing the annotation of DextraDemixer’s base model to its clonotype extension. The box plots indicate the data quartiles with the whiskers extending to the full distribution, excluding outliers outside the 1.5 interquartile range, while the median is indicated as a horizontal line. **g**, Identified CDR3β motifs for LTD and YLQ. Germline-encoded elements of the sequence are highlighted in yellow. Precision of the associated queries to the VDJdb for **h**, LTD, and **i**, YLQ.

First, we applied DextraDemixer’s cell-level prediction to analyze the clonal purity of its annotations. The model identified 370 out of 53,280 and 121 out of 41,565 cells as antigen-specific for the two immunodominant epitopes LTDEMIAQY/HLA-A*01:01 (LTD) and YLQPRTFLL/HLA-A*02:01 (YLQ), respectively (**Fig. 4a**). Because the specificity for a majority of TCRs lacked external experimental validation, we investigated the coherence of the cell-level predicted annotations within the dataset. The pairwise TCRdist^27^ distances between cells assigned to the same specificity were significantly lower than between predicted binder and non-binder cells (**Supp. Fig. 5a**). However, differences between average absolute distances were low (d_LTD_=5.5, d_YLQ_=60.6) due to the presence of multiple distinct sequence clusters within specificity groups. Therefore, we calculated whether its nearest neighbor in sequence space had the same antigen specificity for each TCR. As a random baseline, the frequency of LTD- and YLQ-specific cells in the dataset was 0.7% and 0.3%, respectively. In contrast, the proportion of nearest-neighbor pairs sharing the same specificity annotation was 27.5% for LTD and 68.9% for YLQ (**Fig. 4b**), indicating higher sequence similarity within annotation groups and, thereby, a higher likelihood of the same antigen specificity.

Besides increased sequence similarity, TCRs annotated as epitope-specific by DextraDemixer showed significantly higher *in silico* predicted binding scores derived from TULIP-TCR^28^ for the respective epitope compared to the remaining cells (**Fig. 4c**). We note, however, that the performance of computational binding prediction using TULIP-TCR for LTD is limited (AUC = 0.62), whereas predictions for YLQ are substantially more reliable (AUC = 0.95), as demonstrated in our previous benchmark study^29^. This discrepancy likely explains the minimal shift in prediction scores for LTD-specific TCRs compared to the more pronounced changes observed for YLQ. As an additional internal evaluation, we investigated the coherence of the annotation within cells of the same TCR clonotype as a biological determinant of epitope-specificity. Across all expanded clonotypes, 99.9% received the same annotation. Among clones containing at least one antigen-specific cell, this purity was 87.3% (**Fig. 4d**), indicating that the annotation remained largely coherent not only for non-binding but also for binding clonotypes. We next validated these cell-level predictions by assessing the annotation ambiguity within cells constituting a clonotype. Clones in which 25-75% of cells were labeled epitope-specific were considered ambiguous. Under this definition, only 11.0% of clones and 2.3% of cells belonging to clones with at least one antigen-specific cell could not be assigned a clear specificity (**Fig. 4e**).

To measure the performance of DextraDemixer on this dataset, we defined the TCR-epitope combinations annotated in the original study as positive pairs (*n*_*LTD*_ *= 392*; *n*_*YLQ*_ *= 98*) and supplemented them with TCRs retrieved from VDJdb^26^ that had matching specificities (*n*_*LTD*_ *= 79*; *n*_*YLQ*_ *= 37*). We labeled TCR clones as negative if they were annotated with a different antigen specificity in either the original study (*n*_*LTD*_ *= 1,158*; *n*_*YLQ*_ *= 1,102*) or in VDJdb (*n*_*LTD*_ *= 2,190*; *n*_*YLQ*_ *= 1,632*), as the likelihood of cross-reactivity is considered low. For this evaluation, cells without any annotation or dextramer staining were discarded, resulting in 396 positive and 3,179 negative cells for LTD and 105 and 2,580 such cells for YLQ. Across both epitopes and all three sequencing experiments, DextraDemixer’s base model achieved high separation scores at an average F1-Score of 0.93 (95% CI: 0.85-1.0), and APS score of 0.97 (95% CI: 0.94-1.0) (**Fig. 4f, Supp. Table 3**). Overall, the cell-level predictions exhibited high sequence similarity, *in silico* prediction scores, and high clonal purity.

Expanding the analysis to the clone-aggregated model further led to a small, non-significant increase for both metric scores (Δ_F1_=0.02, Δ_APS_=0.01, **Supp. Fig. 5b**). Lastly, we examined whether DextraDemixer’s clone-aggregated annotations could facilitate the discovery of sequence motifs enriched in antigen-specific TCRs. In LTD-specific CDR3β sequences, the consecutive 3-mer motifs ‘WPY’ and ‘GWP’ were present in 6.5% and 5.6% of sequences, respectively (**Fig. 4g**). While the motifs were not directly germline-encoded, all TCRs containing them were formed by TRBJ2-7 and TRBV29-1. However, this VJ-gene combination alone was not sufficient to identify LTD-specific TCRs in the VDJdb, despite the precision of 2.0% exceeding the database background frequency of 0.02% (**Fig. 4h**). Adding the identified motifs progressively improved precision, ultimately reaching 100% when combined with the VJ genes. In YLQ-specific CDR3β sequences, at least one of the consecutive motifs ‘QNT/ANT’, ‘NTG’, ‘TGE’, and ‘GEL’ encoded by TRBJ2-2 was found in 48 out of 65 DextraDemixer-annotated TCRs (**Fig. 4g**). As with LTD, querying by the motifs identified for YLQ-positive TCRs increased precision relative to the database background frequency and identified 571 out of 2,086 TCRs (27.4%) in VDJdb with the full ‘NTGEL’ motif (**Fig. 4i**). These results demonstrate that DextraDemixer’s specificity annotations can be leveraged to uncover sequence motifs that generalize beyond the analyzed dataset and are associated with shared antigen recognition.

In summary, DextraDemixer proved capable of identifying antigen-specific cells in pMHC-multimer assay data from the SARS-CoV-2 vaccine study. The annotations also agreed closely with external validation data and supported the identification of antigen-specific motifs.

## Discussion

In summary, we presented DextraDemixer, a hierarchical Bayesian mixture model to robustly identify antigen-specific T cells in pMHC-multimer experiments. We evaluated it across simulated and multiple real-world datasets spanning different experimental platforms, including 10x Genomics and BD Rhapsody. DextraDemixer demonstrated consistent and accurate classification despite substantial differences in sequencing depth and UMI count distributions. In contrast to existing approaches, DextraDemixer enables explicit control of the FDR, thereby improving the reliability of downstream analyses, including motif discovery, cluster-based specificity imputation, and the generation of training data for machine learning models. Such applications are particularly sensitive to false-positive antigen assignments, as they obscure true sequence patterns, dilute specificity-associated motifs, and introduce misleading labels into TCR-specificity training data, a key issue that plagues current large-scale data collection efforts^30,31^.

While DextraDemixer provides accurate classification, several avenues for model refinement remain. First, the model could be extended to incorporate sequence similarity between TCRs, reflecting the biological observation that similar TCRs often share antigen specificity^27^, rather than relying solely on exact clonotype matches. This extension becomes particularly important when adapting DextraDemixer to B-cell specificity assays, where somatic hypermutations frequently alter receptor sequences while largely preserving the original antigen specificity. Second, DextraDemixer assumes independence between pMHCs, relaxing this assumption to explicitly model interactions, such as competitive binding or cross-reactivity, could further improve inference and enable more accurate calibration of background noise. Third, although stochastic variational inference provides a computationally efficient approximation of the posterior distribution, a more precise estimation could be achieved with Markov Chain Monte Carlo sampling through highly parallelized computation.

Importantly, while DextraDemixer accurately infers TCR-pMHC binding, it is crucial to note that physical interaction, though a necessary component of antigen recognition, does not by itself guarantee functional responsiveness. The immunological consequence of recognition depends on additional factors, including TCR affinity and avidity, antigen density, co-stimulation, as well as differentiation and cellular state of the T cell. Thus, a T cell may show detectable pMHC-multimer binding without mounting effector responses. Therefore, DextraDemixer-derived probabilities should be interpreted as uncertainty-aware antigen-specificity assignments and ideally integrated with orthogonal readouts such as transcriptomic profiles, cytokine measurements, or other functional assays to obtain a more complete view of the immune response.

Ultimately, DextraDemixer provides an accurate, statistically grounded, and fully automated framework to deconvolve biological signals from noise in pMHC-multimer sequencing experiments. By replacing manually tuned thresholds and dataset-specific heuristics with uncertainty-aware inference and FDR-calibrated classification, the method supports more scalable, reproducible, and standardized analyses across users, datasets, platforms, and experimental conditions. Its computational scalability and integration with the scverse^32^ ecosystem further make DextraDemixer readily accessible for routine use in immunological research.

## Methods

### Model description

DextraDemixer aims to distinguish antigen-specific binding T cells from non-specific binders and assay noise based on measured pMHC-multimer UMI counts. Let *X* ∈ ℕ^*N*×*M*^ denote the count matrix of *N* cells measured against *M* antigens, where cells are grouped into clonotypes *c* ∈ {1, …, *C*} that share the same TCR sequence. To separate antigen-specific signal from background, we employ a fully Bayesian hierarchical negative binomial (NB) mixture model. The model is formulated under the assumptions that (1) each antigen’s measured count distribution *X*_.*j*_ is independent and identically distributed, (2) the counts follow a mixture of two NB distributions representing signal (antigen-specific binder) and noise (technical noise, non-specific binder), respectively, (3) all cells of a clonotype *c* belong either to the antigen-specific signal or noise component, (4) cells from the noise component exhibit a lower expected read count than specifically binding cells.

As mixture models are highly susceptible to poor fits due to many local minima, DextraDemixer utilizes a data-informed initialization strategy to guide the hyperpriors. Specifically, k-means clustering with *k* = 2 is performed, and the resulting cluster means 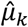, variance 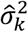, and proportions 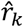 are used to calibrate the prior distributions. To mitigate the effect of varying sequencing depth for each cell, size factor normalization values ŝ_*i*_ are calculated following DESeq2^33^.

Based on the laid out assumptions, we derived the following hierarchical model considering each antigen *j* individually (in the following, *j* is dropped for notational simplicity):

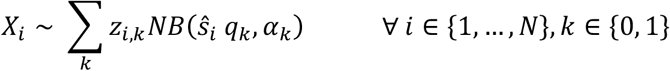

Here, the NB distribution is parameterized by its mean *q*_*k*_, and inverse dispersion *α*_*k*_ for the signal (*k* = 1) and noise (*k* = 0) component individually. The one-hot encoded latent class variable *z*_*i*_ is drawn from a categorical distribution (Cat) with its mixture weights *π* following a Dirichlet (Dir) distribution with prior parameters determined by the k-means cluster ratios 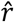:

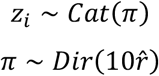

The component means *q*_*k*_ are assigned Log-Normal (LogN) prior distributions, with their parameters 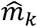 and 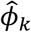 calculated to match the observed mean and variance of their respective k-means clusters. An inequality constraint is imposed on *q*_1_ and *q*_0_ following assumption (4) to guarantee identifiability:

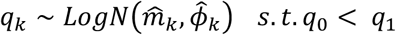

To set the NB’s variance in relationship to its mean, we draw the coefficient of dispersion *τ*_*k*_ from the non-informative Half-Chauchy (HC) prior distribution. This imposes a linear mean-variance relationship, which restricts the signal component from erroneously expanding its variance to fit uncaptured probability mass from the noise distribution.

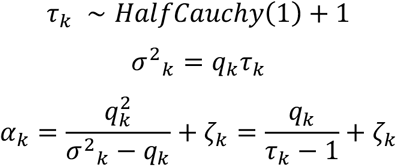

To further regularize the signal component against absorbing residual background mass, particularly in datasets exhibiting extreme class imbalance and low signal-to-noise ratio, we add a deterministic offset *ζ*_*k*_ set to *ζ*_1_ = 5, *ζ*_0_ = 0 to the inverse dispersion parameter *α*_*k*_.

#### Negative controls

To approximate assay noise and unspecific binding, negative control samples *X*_*i*_^*neg*^ (usually an identical HLA allele loaded with an irrelevant epitope^34^ or a HLA-mismatched pMHC-complex^22^) are often included alongside the pMHCs of interest. If available, DextraDemixer jointly models those to calibrate the noise component of the hierarchical model. As we observed distributional shifts in the mean and dispersion between the negative control and noise component in the pMHC data from 10x Genomics^31^ (**Supp. Fig. 2a**), we extended DextraDemixer to model this relationship as inverse scale factors for mean *s*_*q*_ and dispersion *s*_*τ*_ drawn from log-normal priors fitted on these data:

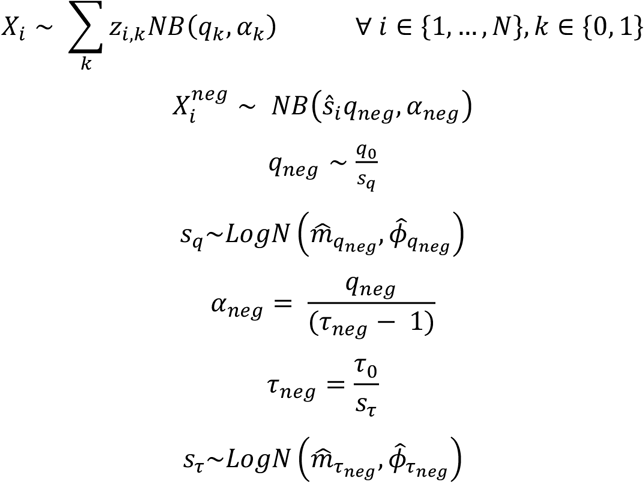

#### Inference

To infer the model parameters, we employed Blackbox Variational Inference (BBVI)^35^ with an Auto-Normal Guide. Because BBVI cannot directly differentiate through discrete variables, we marginalize the latent class ***z*** and reconstruct it through the obtained posterior class probabilities *p*_*i*_ for each sample:

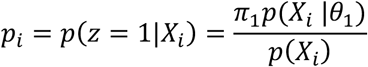

BBVI is performed over 1,000 iterations using gradient-clipped Adam^36^ (clip=10). The initial learning rate is set to 0.3 and exponentially decayed with a decay rate of 0.995, reaching a final learning rate of 0.003. Model parameters are initialized using the set with the lowest loss out of 10 random initializations, based on the medians of the model’s priors. Finally, to leverage the biological premise that cells expressing identical TCR share the same antigen specificity, we aggregate the individual cell-level posteriors to compute the median assignment probability *p*_*i*_^*c*^ for each clonotype with interpolation set to the higher value.

#### Balanced and dynamic thresholding with control of false discovery rate

To assign cells a discrete signal or noise label, we evaluated the expected posterior class probability on a cell-*p*_*i*_ or clone-median *p*_*i*_^*c*^ level against a threshold *t*. As a balanced decision boundary between Type I and Type II errors, a threshold *t* = 0.5 was used as the default method. However, as established in earlier works^37,38^, Bayesian models need to be controlled for multiplicity when making decisions, to prevent an inflation of false positives. To this end, we employ the direct posterior probability approach to control the false discovery rate^39^. Briefly, if 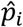 approximates the probability of a cell *i* belonging to the signal component, then the local FDR, probability of a Type I error, is defined as 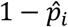. For a threshold *t*, we then obtain a set of predicted antigen-specific cells 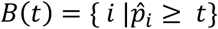. The global false discovery rate 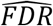 for a threshold *t* is then approximately given by:

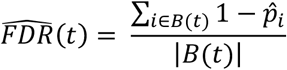

For a user-specific target FDR *α*, we find the optimal threshold *t*’ that includes as many cells as possible whose approximated global FDR does not exceed the target FDR *α*.

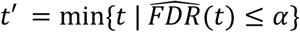

We extended this framework to leverage the full posterior distribution. Rather than relying solely on posterior expectations, we computed the posterior distribution of the global FDR across all inference draws for each candidate threshold. We then apply a credible interval-bounded control strategy, selecting the minimal threshold *t*′ that ensures the posterior probability of the global FDR remaining below the target *α* meets a predefined credibility level *γ* of 0.5:

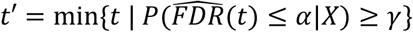

We finally obtain the set of credibly binding cells *B*(*t*’). The same holds for clonotype-aggregated probabilities. Once credible binding clonotypes have been determined, all cells of these clonotypes will be deemed credible binding cells.

### Simulation

We developed a simulation framework (**Supp. Fig. 1**) to generate datasets with a wide range of scenarios to comprehensively benchmark, evaluate, and analyze our model’s robustness. To ensure faithfulness, we determined distribution parameters and their relationships on experimental human PBMC and murine spleen single-cell pMHC-multimer data from 10x Genomics^40^ (“10k Human A0201 PBMCs with CMV, Flu, and SARS-Cov2 Spike In”; “5k Human A0201 | B0702 PBMCs”; “2k Mouse H2Kb OT-1 Splenocytes”; “10k Human A2402 PBMCs with EBV and CMV Spike In”) (**Supp. Fig. 2**). To isolate signal and noise distributions, a threshold based on the maximum outlier-filtered (<0.999 quantile) negative control count was applied. pMHCs for which both components had more than 20 samples were used for further analysis. Maximum likelihood estimation was subsequently applied to negative control, signal, and noise UMI counts to fit negative binomial distributions, yielding empirical means and variances (**Supp. Fig. 2b,c**). The distribution of component means was then approximated with a truncated log-normal distribution and fitted using a weighted Method of Moments (**Supp. Fig. 2d,e**). Because empirical data exhibited a strong log-linear relationship between mean and variance (**Supp. Fig. 2f**), therefore we modeled the variance conditionally on the mean and added a Gaussian deviation term to capture the residual error of the fit (**Supp. Fig. 2g**). Finally, the clonal size distribution was fitted by a Boltzmann distribution to “10k Human A0201 PBMCs with CMV, Flu, and SARS-CoV-2 Spike In”, as the other datasets exhibited unrealistic and artificial clone size distributions driven by arbitrary spike-in ratios (**Supp. Fig. 2h**).

Synthetic datasets were parameterized by the following variables: total cell count *N*, total clonotypes *C*, binding probability *p*, probability of clonal dropout *p*_*out*_, and signal-to-noise ratio *β*. We generated a cell *i* with its clone *c*, pMHC UMI count data *X*_*i*_, and a negative control *X*_*i*_^*neg*^ as follows (**Supp. Fig. 1**): First, we simulated the *C* clone sizes and assigned one cell to each clone to prevent empty classes. The remaining *N* − *C* leftover cells were then distributed with a Multinomial distribution parameterized by sampling from the previously fitted Boltzmann distribution, obtaining the final number of cells per clonotype *n*_*c*_. The binding class *z*_*c*_ for each clonotype was drawn from a Bernoulli distribution with *p* as the probability belonging to the binder class *z*_*c*_ = 1. To enforce the target ratio *p*, we sampled 10,000 times and selected the trial with the lowest absolute error to the target on a cell- and clone-level. If the error exceeded a tolerance of 5%, the sampling framework restarted with a newly sampled set of clone sizes. Next, we generated the pMHC UMI count *X*_*i*_ per cell *i* given their clone’s class assignment. We sampled the noise component mean *q*_0_ from the fitted truncated Log-Normal and set the signal component mean as *q*_1_ = *β* ⋅ *q*_0_ with the user-defined signal-to-noise ratio *β*. The corresponding variances 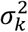 were computed with the fitted log-linear model, augmented with a sampled Gaussian residual deviation *ϵ*_*k*_. The Negative Binomial inverse dispersion parameter was then derived as 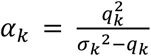. Then, we sampled *n*_*c*_ times from an NB with corresponding parameters dictated by *z*_*c*_ for each clonotype *c*. To further introduce intrinsic variance, each noise clone’s inverse dispersion value got perturbed with a Gaussian as 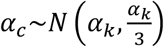. If dropout cells of antigen-specific clones were simulated, a dropout state vector 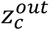 of length *n*_*c*_ was sampled from a Bernoulli with probability *p*_*out*_. Cells designated as dropouts were assigned UMI counts drawn from the noise instead of the signal distribution. Negative control read counts for each cell *i* were similarly simulated, first sampling *q*_*neg*_ from the specifically fitted truncated Log-Normal, then estimating 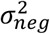 and calculating *α*_*neg*_ accordingly. Different random seeds were used in each repetition to produce different datasets with the same hyperparameter sets.

For the synthetic performance benchmark, we used the following parameters:

Repetitions: 5

*N*: {100, 200, 500, 1000, 2000, 5000, 10000}

*C*: 0.4*N*

*p*: {0.001, 0.002, 0.005, 0.01, 0.02, 0.05, 0.1, 0.2, 0.5 | *N* ⋅ *p* ≥ 1}

*p*_*out*_: {0.0}

*β*: {10, 15, 20, 30, 40, 50, 100, 200}

For the clone outlier benchmark, we used the following parameters: Repetitions: 5

*N*: {1,000, 10,000}

*C*: 0.4*N*

*p*: {0.1}

*p*_*out*_: {0.0, 0.05, 0.1, 0.15, 0.2, 0.25, 0.3, 0.35, 0.4, 0.45,

0.5, 0.55, 0.6, 0.65, 0.7, 0.75, 0.8, 0.85, 0.9, 0.95}

*β*: {20, 50}

For the runtime benchmark, we used the following parameters:

Repetitions: 5

*N*: {10000, 20000, 40000, 60000, 80000, 100000}

*C*: 0.4*N*

*p*: {0.01, 0.1}

*p*_*out*_: {0.0}

*β*: {20, 50}

### Analysis of publicly available datasets

#### Single-cell data of the CMV-positive spike-in experiment by Gemünd et al

We downloaded the single-cell immune profiling and pMHC multimer dataset of Gemünd et al.^25^ alongside the original preprocessing and analysis script from the associated GitLab repository (https://gitlab.dzne.de/ag-beyer/gemuend_cmv_2025). Briefly, the dataset consists of two sets of epitope-specific, polyclonal T cells (NLVPMVATV - pp65_495-503_ presented on HLA-A∗02:01, CRVLCCYVL - IE-1_309-317_ on HLA-C∗07:02) and one set of T cells enriched for TPRVTGGGAM-specificity (pp65_417-426_ on HLA-B∗07:02), which were spiked-in (as 1:1:1-mix) into a HLA-matched and mismatched background samples at two different concentrations (at 5% and 50% against the two backgrounds (BG), respectively). BD-Rhapsody-based single-cell immune profiling and pMHC dextramer sequencing for three dextramers targeting the respective CMV epitopes: DexA (HLA-A∗02:01/pp65495-503), DexB (HLA-B∗07:02/pp65417-426), and DexC (HLA-C∗07:02/IE-1309-317), as well as a negative control DexN (HLA-B∗08:01/AAKGRGAAL), for a total of 10,000 loaded T cells was performed. The single-cell data was processed according to the original scripts. For evaluation, four sets were created: DexA 5%, DexA 50%, DexC 5%, and DexC 50%. Spike-in clones recognizing the respective DexA or DexC dextramer were labeled as positive, whereas T cells derived from the HLA-mismatched background, as well as the spike-in clones binding to other epitopes were labeled as negatives. Further, each set was split into a setting using only cells with complete TCR sequences, and another using all cells, where we treated cells without a full TCR as belonging to a clonotype of a single cell.

We used this setup to quantitatively compare the full DextraDemixer against BEAM^23^, ICON^21^, and ITRAP^22^. For ICON, we implemented a version of the code reproducing their github-repository algorithm and a version reproducing the methods as described in the manuscript. In the main figure, we show the results for the manuscript version, which had the better performance (**Supp. Fig. 4**). For ITRAP, a method based on filtering out erroneous signals from the dataset based on a set of diverse criteria, we only applied filters regarding the dextramer signal (i.e., optimal threshold, and specificity multiplets), since the dataset was already filtered for incomplete TCRs, non-viable cells, and hashing duplicates using the original authors script.

To evaluate whether DextraDemixer identifies antigen-specific activation signatures, we compared CD8+ TPR-enriched cells predicted as DexB^+^ or DexB^-^ by DextraDemixer. To analyze this in an unbiased manner, we used the antigen-specific T cell activation gene programs systematically identified in a large-scale T cell atlas comprising 1.7 million T cells from 38 human tissues and five disease contexts^24^. We further identified predicted specificities of the cells by querying the receptors to VDJdb^26^ following the original publication using scirpy^41^. In particular, a receptor was considered to match a receptor from VDJdb when there was complete identity in the amino acid chain of any of the receptor arms in either the primary or secondary chain.

#### Single-cell data of the SARS-CoV-2 vaccine cohort by Kocher et al

The single-cell immune profiling data and experimental verifications of antigen-specific clones were collected in the context of our original publication^11^.

DextraDemixer with cell-level and clone-aggregated predictions were applied to LTD, and YLQ UMI counts for all cells stained with the corresponding pMHC multimers. The original study did not include negative control multimers, and only a limited subset of cells was evaluated with multiple pMHC. Consequently, we omitted the joint modelling of negative controls and size factor normalization steps, respectively. Pairwise distances were calculated for all clones in the dataset using TCRdist3^42^. TULIP-TCR^28^ binding scores were predicted for both epitopes through the ePytope-TCR interface^29^. Clonal purity was defined as the fraction of cells that have the specificity annotation of the majority class within this clone. Classification metrics were calculated per sequencing experiment based on the positive or negative labels described above, filtering all cells without any label. VDJdb^26^ (version 2025-02-21) was queried for exact CDR3β sequence identity towards other epitopes to define negative labels. To identify antigen-specific motifs, all 3-mers from the CDR3β sequences of the dataset were formed, and their occurrence was counted for antigen-specific and all other clonotypes. Statistical motif enrichment was determined by Fisher’s exact test on these counts and the number of clones in each category at a significant threshold of *p<0*.*05*. The remaining analysis was conducted on all motifs occurring more than five times in the epitope-specific sequences that exhibit at least a ten-fold higher fractional frequency in epitope-specific compared to unspecific sequences.

## Data availability

The synthetic datasets have been deposited to Zenodo at https://doi.org/10.5281/zenodo.20759653. The single-cell datasets can be found in their respective public repositories: The processed data and source code to analyze the sequencing data of the experiment by Gemünd et al. can be accessed on GitLab at https://gitlab.dzne.de/ag-beyer/gemuend_cmv_2025. The SARS-CoV-2 vaccine cohort data by Kocher et al. can be found at NCBI GEO under the accession numbers GSE261966, GSE261967, and GSE249998. Processed and annotated data can be accessed at Zenodo (https://doi.org/10.5281/zenodo.13981507). The source code to analyze the sequencing data is available on GitHub at https://github.com/SchubertLab/CovidVac_CD8.

## Code availability

The method has been implemented in Python 3.13 using Numpyro 0.19, numpy 2.3.5, jax 0.8.1, muon 0.1.7, and scirpy 0.22.3. Source code has been deposited on GitHub at https://github.com/SchubertLab/DextraDemixer. All code and results to reproduce the presented analyses can be found on GitHub at https://github.com/SchubertLab/DextraDemixer_reproducibility. All tested methods have been integrated into a unifying Python API that can directly interact with Scirpy and muData.

## Acknowledgement

We thank Ioanna Gemünd for our valuable discussion on usability requirements for the software package. During the preparation of this work, the authors used Gemini by Google to improve the readability and conciseness of the manuscript. After using this tool/service, the authors reviewed and edited the content as needed and take full responsibility for the content of the publication.

## Author Contributions

The conception and design of the study was done by BS.

Model development was done by YA, BS, IB.

The data were acquired by KS, MG.

Data analysis and interpretation were performed by YA, IB, FD.

The manuscript was written by YA, BS, IB, FD.

All authors read and reviewed the manuscript.

## References

1. Hennecke, J. & Wiley, D. C. T cell receptor-MHC interactions up close. Cell 104, 1–4 (2001).

2. Boutet, S. C. et al. Scalable and comprehensive characterization of antigen-specific CD8 T cells using multi-omics single cell analysis. J Immunol 202, 131. 4–131.4 (2019).

3. Ma, K.-Y. et al. High-throughput and high-dimensional single-cell analysis of antigen-specific CD8 T cells. Nat Immunol 22, 1590–1598 (2021).

4. Ulbrich, J., Lopez-Salmeron, V. & Gerrard, I. BD Rhapsody™ Single-Cell Analysis System Workflow: From Sample to Multimodal Single-Cell Sequencing Data. Methods Mol Biol 2584, 29–56 (2023).

5. Ng, A. H. C. et al. MATE-Seq: microfluidic antigen-TCR engagement sequencing. Lab Chip 19, 3011–3021 (2019).

6. Zhang, S.-Q. et al. High-throughput determination of the antigen specificities of T cell receptors in single cells. Nat. Biotechnol. 36, 1156–1159 (2018).

7. Luo, S., Notaro, A. & Lin, L. ATLAS-seq: a microfluidic single-cell TCR screen for antigen-reactive TCRs. Nat Commun 16, 216 (2025).

8. Adams, B. A. et al. An integrated reagent and multimodal analysis workflow to enrich and characterize peptide-specific CD8+T cells. J Immunol 210, 249. 17–249.17 (2023).

9. Zhang, R. et al. Whole-protein screening and multi-modal profiling of antigen-specific CD4+ T cells at single-cell resolution. Nat. Commun. 17, 3979 (2026).

10. Kocher, K. et al. Adaptive immune responses are larger and functionally preserved in a hypervaccinated individual. Lancet Infect. Dis. 24, e272–e274 (2024).

11. Kocher, K. et al. Vaccination-induced T cell responses maintain polyclonality with high antigen receptor avidity. Sci Immunol 10, eadu6730 (2025).

12. Inekci, D., Hansen, B. E. & Brix, L. Simultaneous detection and characterization of antigen-specific B cells and CD4+ and CD8+ T cell responses upon natural infection and vaccination. J. Immunol. 210, 251. 10–251.10 (2023).

13. Frischholz, S. et al. Metabolic quiescence of naive-like memory T cells precedes and maintains antigen-specific T cell memory. Nat. Immunol. 27, 452–462 (2026).

14. Minervina, A. A. et al. SARS-CoV-2 antigen exposure history shapes phenotypes and specificity of memory CD8+ T cells. Nat. Immunol. 23, 781–790 (2022).

15. Adamo, S. et al. Signature of long-lived memory CD8+ T cells in acute SARS-CoV-2 infection. Nature 602, 148–155 (2022).

16. Lenkala, D. et al. NEO-STIM advances personalized neoantigen-specific adoptive T cell therapy. Nat. Commun. 17, 3683 (2026).

17. Eggebø, M. S. et al. TCR-engineered T cells targeting a shared β-catenin mutation eradicate solid tumors. Nat. Immunol. 26, 1726–1736 (2025).

18. Salvatori, E. et al. Neoantigen cancer vaccine augments anti-CTLA-4 efficacy. NPJ Vaccines 7, 15 (2022).

19. 10x Genomics. A New Way of Exploring Immunity - Linking Highly Multiplexed Antigen Recognition to Immune Repertoire and Phenotype - 10x Genomics. 10x Genomics https://www.10xgenomics.com/resources/application-notes/a-new-way-of-exploring-immunity-linking-highly-multiplexed-antigen-recognition-to-immune-repertoire-and-phenotype/ (2019).

20. Francis, J. M. et al. Allelic variation in class I HLA determines CD8+ T cell repertoire shape and cross-reactive memory responses to SARS-CoV-2. Sci. Immunol. 7, eabk3070 (2022).

21. Zhang, W. et al. A framework for highly multiplexed dextramer mapping and prediction of T cell receptor sequences to antigen specificity. Sci. Adv. 7, eabf5835 (2021).

22. Povlsen, H. R. et al. Improved T cell receptor antigen pairing through data-driven filtering of sequencing information from single cells. Elife 12, (2023).

23. 10x Genomics. Cell Ranger’s Antigen Capture (BEAM) Algorithm. https://www.10xgenomics.com/support/software/cell-ranger/latest/algorithms-overview/cr-5p-antigen-algorithm.

24. Kotliar, D. et al. Reproducible single-cell annotation of programs underlying T cell subsets, activation states and functions. Nat. Methods 22, 1964–1980 (2025).

25. Gemünd, I. et al. Characterization of human CMV-specific CD8+ T cells using multi-layer single-cell omics. Cell Rep. Methods 5, 101085 (2025).

26. Goncharov, M. et al. VDJdb in the pandemic era: a compendium of T cell receptors specific for SARS-CoV-2. Nat. Methods 19, 1017–1019 (2022).

27. Dash, P. et al. Quantifiable predictive features define epitope-specific T cell receptor repertoires. Nature 547, 89–93 (2017).

28. Meynard-Piganeau, B., Feinauer, C., Weigt, M., Walczak, A. M. & Mora, T. TULIP: A transformer-based unsupervised language model for interacting peptides and T cell receptors that generalizes to unseen epitopes. Proc. Natl. Acad. Sci. U. S. A. 121, e2316401121 (2024).

29. Drost, F. et al. Benchmarking of T cell receptor-epitope predictors with ePytope-TCR. Cell Genom. 5, 100946 (2025).

30. Messemaker, M. et al. A functionally validated TCR-pMHC database for TCR specificity model development. bioRxivorg (2025) doi:10.1101/2025.04.28.651095.

31. Jouannet, C., Vantomme, H., Gouge, K. L., Klatzmann, D. & Mariotti-Ferrandiz, E. Benchmarking unsupervised methods for inferring TCR specificity. NAR Genom. Bioinform. 7, lqaf150 (2025).

32. Virshup, I. et al. The scverse project provides a computational ecosystem for single-cell omics data analysis. Nat. Biotechnol. 41, 604–606 (2023).

33. Love, M. I., Huber, W. & Anders, S. Moderated estimation of fold change and dispersion for RNA-seq data with DESeq2. Genome Biol. 15, 550 (2014).

34. Newell, E. W., Klein, L. O., Yu, W. & Davis, M. M. Simultaneous detection of many T-cell specificities using combinatorial tetramer staining. Nat. Methods 6, 497–499 (2009).

35. Ranganath, R., Gerrish, S. & Blei, D. M. Black Box Variational Inference. arXiv [stat.ML] 814–822 (2013).

36. Kingma, D. P. & Ba, J. Adam: A method for stochastic optimization. arXiv [cs.LG] (2014).

37. Müller, P., Parmigiani, G. & Rice, K. FDR and Bayesian multiple comparisons rules. In Bayesian Statistics 8 359–380 (Oxford University PressOxford, 2007).

38. Scott, J. G. & Berger, J. O. Bayes and empirical-Bayes multiplicity adjustment in the variable-selection problem. aos 38, 2587–2619 (2010).

39. Newton, M. A., Noueiry, A., Sarkar, D. & Ahlquist, P. Detecting differential gene expression with a semiparametric hierarchical mixture method. Biostatistics 5, 155–176 (2004).

40. 10X Genomics. Dataset Collection. https://www.10xgenomics.com/datasets.

41. Sturm, G. et al. Scirpy: a Scanpy extension for analyzing single-cell T-cell receptor-sequencing data. Bioinformatics 36, 4817–4818 (2020).

42. Mayer-Blackwell, K. et al. TCR meta-clonotypes for biomarker discovery with tcrdist3 enabled identification of public, HLA-restricted clusters of SARS-CoV-2 TCRs. Elife 10, (2021).

