## Supplementary Figures and Notes for "DextraDemixer enables accurate identification of antigen-specific T cells from pMHC multimer experiments"

### Supplementary Text

We address the problem of distinguishing true antigen-binding T cells from non-specific and technical noise in pMHC-multimer assays using a fully Bayesian hierarchical negative-binomial mixture model. To this end, we assume that **(1)** each antigen's measured count distribution $X_{.j}$ is independent and identically distributed, **(2)** the counts follow a mixture of two NB distributions representing signal (antigen-specific binder) and noise (technical noise, unspecific-binder), **(3)** all cells of a clonotype $c$ belongs either to the antigen-specific signal or noise component, **(4)** cells from the noise component exhibit a lower expected read count than specifically binding cells. Based on these assumptions, we derive the following hierarchical model, given target pMHC-multimer and optional negative control pMHC-multimer UMI counts:

$$X_i \sim \sum_k z_{i,k} NB(\hat{s}_i\, q_k, \alpha_k) \qquad \forall\, i \in \{1, \ldots, N\}, k \in \{0, 1\}$$

$$z_i \sim Cat(\pi)$$

$$\pi \sim Dir(10\hat{r})$$

$$q_k \sim LogN(m_k, \phi_k) \quad s.t.\, q_0 < q_1$$

$$\tau_k \sim HalfCauchy(1) + 1$$

$$\sigma^2{}_k = q_k \tau_k$$

$$\alpha_k = \frac{q_k^2}{\sigma^2{}_k - q_k} + \zeta_k = \frac{q_k}{\tau_k - 1} + \zeta_k$$

**Jointly modeling negative controls:**

$$X_i^{neg} \sim NB\left(\hat{s}_i q_{neg}, \alpha_{neg}\right)$$

$$q_{neg} \sim \frac{q_0}{s_q}$$

$$s_q \sim LogN\left(\hat{m}_{q_{neg}}, \hat{\phi}_{q_{neg}}\right)$$

$$\alpha_{neg} = \frac{q_{neg}}{(\tau_{neg} - 1)}$$

$$\tau_{neg} = \frac{\tau_0}{s_\tau}$$

$$s_\tau \sim LogN\left(\hat{m}_{\tau_{neg}}, \hat{\phi}_{\tau_{neg}}\right)$$

With $\hat{\mu}_k$, $\hat{\sigma}_k^2$, and $\hat{r}_k$ being the mean, variance, and cluster size ratios of a k-means clustering (k=2), $\hat{s}_i$ a per cell size factor following DESeq2[35]. $\hat{m}_{q_{neg}}, \hat{\phi}_{q_{neg}}$, $\hat{m}_{\tau_{neg}}, \hat{\phi}_{\tau_{neg}}$ are hyperprior parameters derived from the pMHC-multimer data from 10x Genomics[40].

### Supplementary Figures

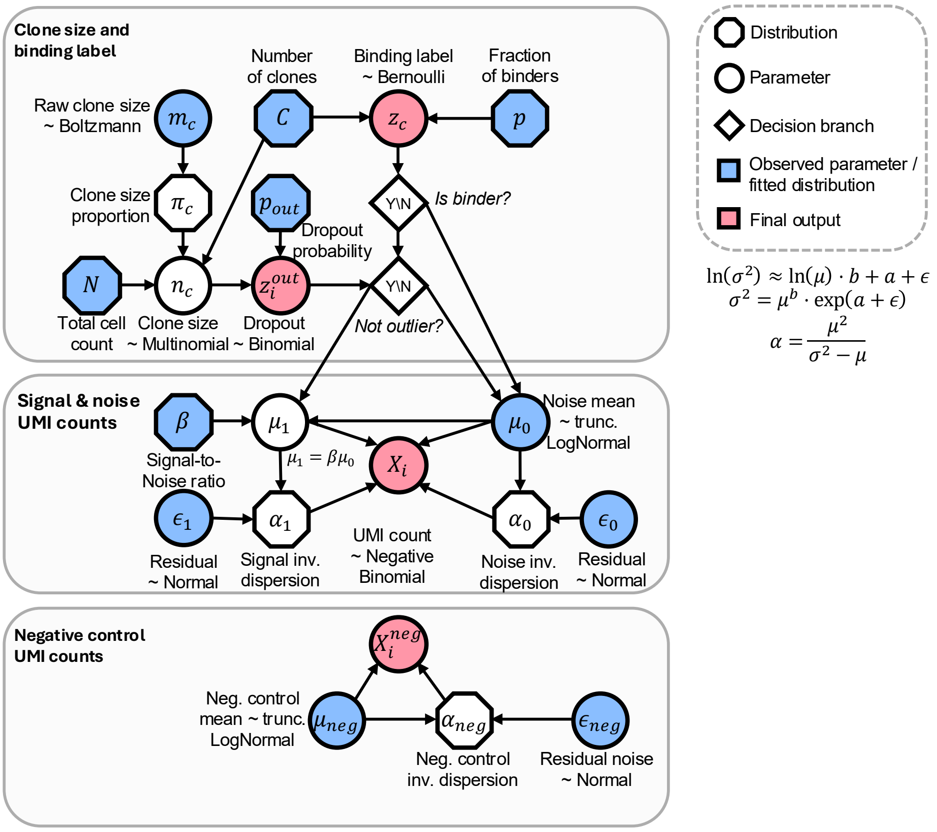


**Supplementary Figure 1 | Graphical model of the simulation framework.** Directed acyclic graph (DAG) describing the generative process for simulating clone-resolved UMI counts with binder status, dropout, and noise components. Blue nodes denote observed or user-specified variables (or sampled from fitted distributions), white nodes denote latent variables and parameters, diamonds indicate decision branches, and red nodes indicate final simulated outputs.

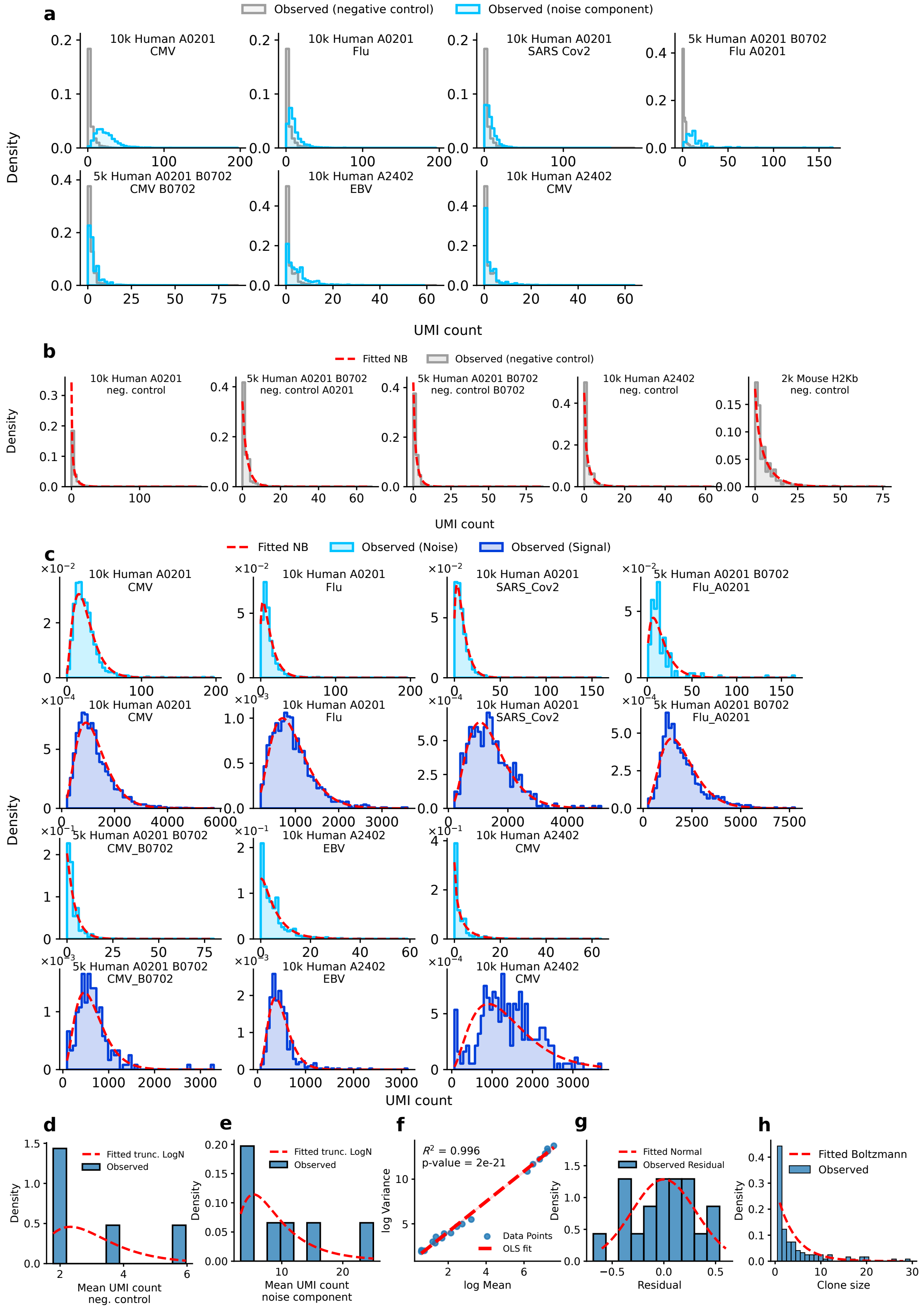


**Supplementary Figure 2 | Estimation of distribution parameters from four 10x Genomics single-cell pMHC datasets. a,** Outlier filtered negative control and noise component. **b, c,** Fitted Negative Binomial (NB) distributions on observed **b,** outlier-removed negative control UMI counts **c,** noise and signal component UMI counts, determined with negative control-based threshold. **d, e,** Fitted truncated log-normal distribution on mean UMI count of observed **d,** negative control, **e,** noise component. **f, g,** log-linear relationship between NB-determined mean and variance of UMI count; **f,** fitted linear regression curve; **g,** residual between modeled and observed relationship. **h,** Observed and Boltzmann-fitted clone size distribution on 10k Human A0201. Full 10x Genomics dataset[31] IDs - "10k Human A0201": "10k Human A0201 PBMCs with CMV, Flu, and SARS-Cov2 Spike In"; "5k Human A0201": "5k Human A0201 B0702": "5k Human A0201 | B0702 PBMCs"; "2k Mouse H2kb": "2k Mouse H2Kb OT-1 Splenocytes"; "10k Human A2402 PBMCs with EBV and CMV Spike In"

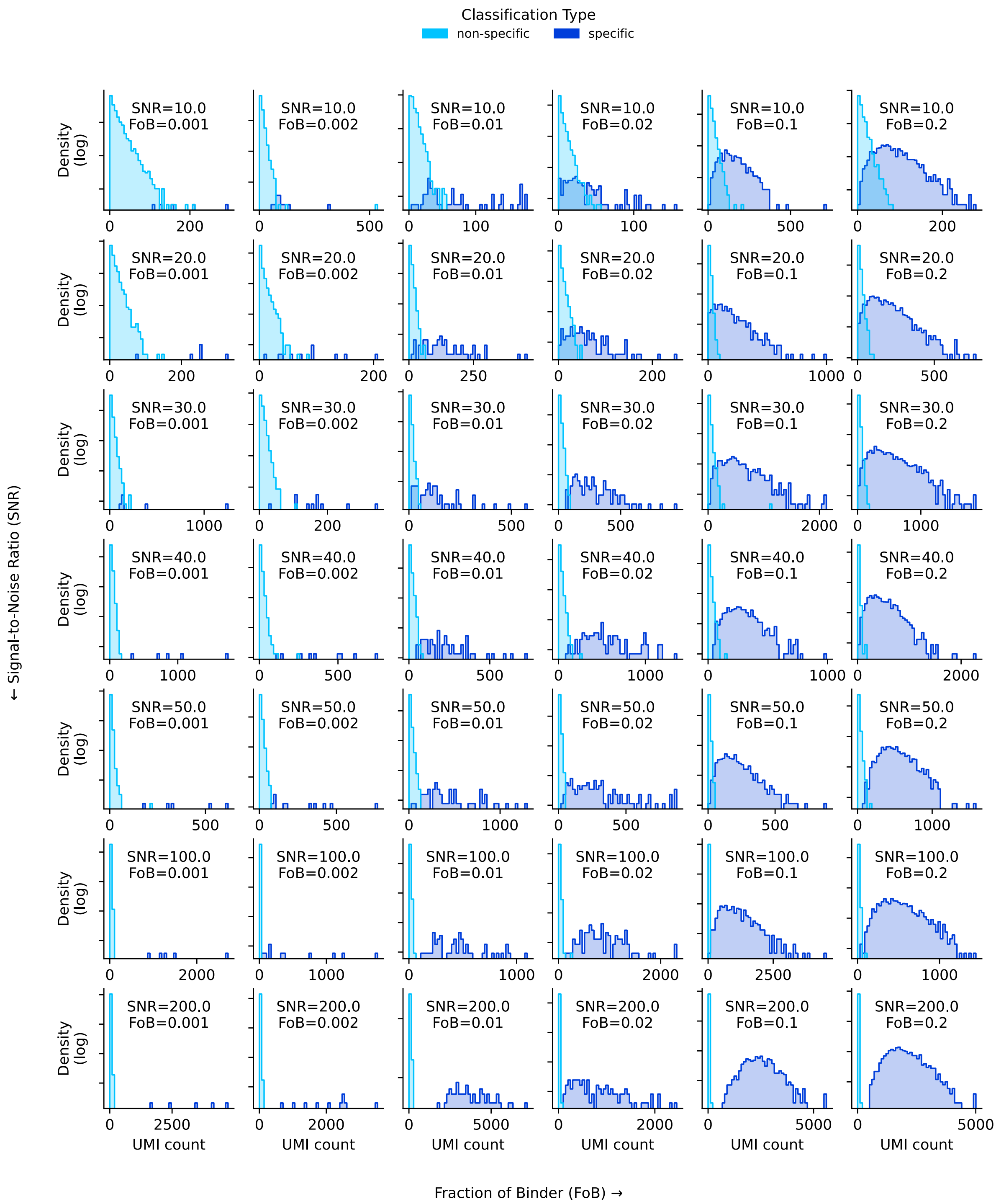


**Supplementary Figure 3 | Log-scaled simulated UMI count distributions.** Signal-to-noise ratios (SNRs) increase with rows, and the fraction of binders (FoB) increases with columns. All shown examples have a total cell count of 5,000. To make an unbiased selection, the first repetition was chosen.

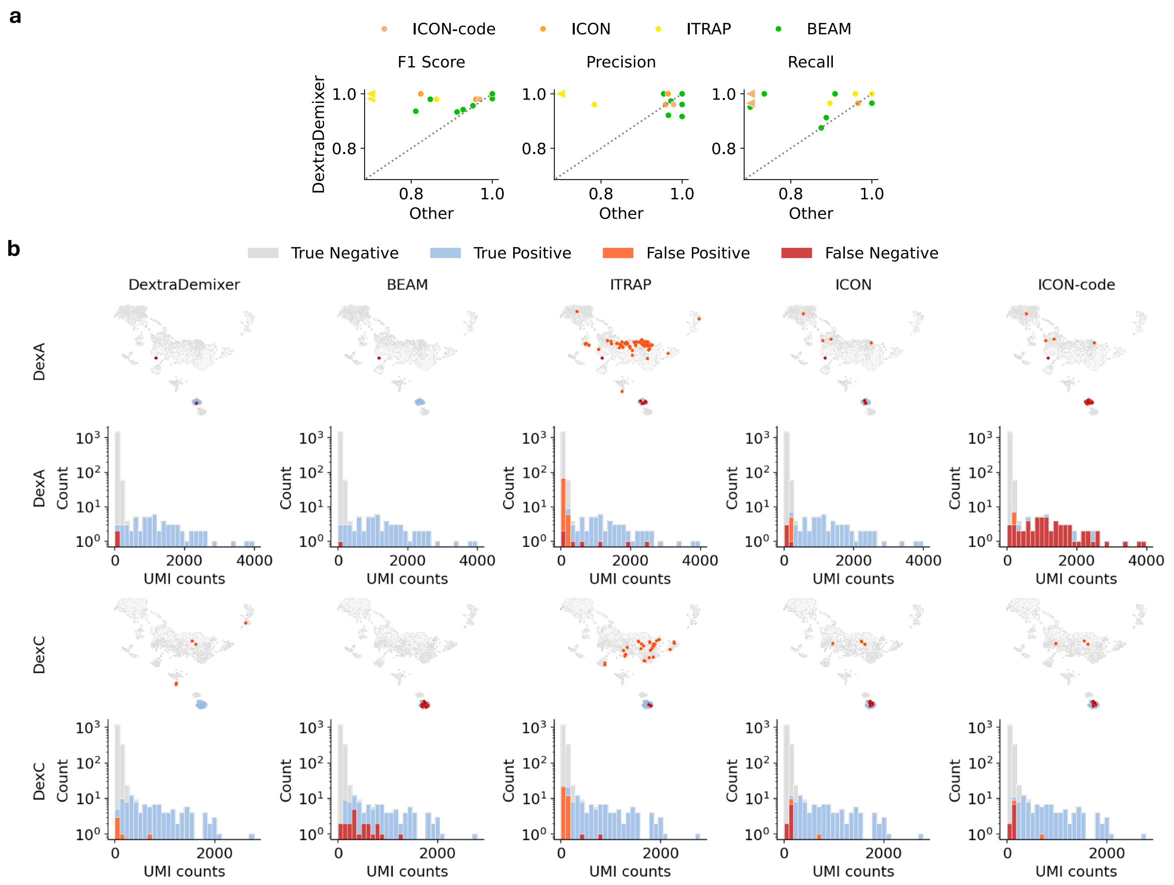


**Supplementary Figure 4 | Pairwise benchmark comparison and classification errors in the clonal spike-in experiment. a,** Pairwise comparison of DextraDemixer against BEAM, ITRAP, ICON and ICON-code across spike-in benchmark settings. Scatter plots show F1 score, precision, and recall, with DextraDemixer on the y axis and the respective comparator method on the x axis. Each point represents one benchmark setting. The diagonal indicates equal performance; points above the diagonal indicate higher performance of DextraDemixer. When performance is lower than 0.8, the method is depicted with an arrow instead of a point. **b,** Method-specific classification results for DexA and DexC spike-in cells visualized on the single-cell UMAP embedding and corresponding dextramer UMI count distributions. Histograms show the distribution of dextramer UMI counts for each method and dextramer, illustrating how classification errors relate to the underlying signal distributions.

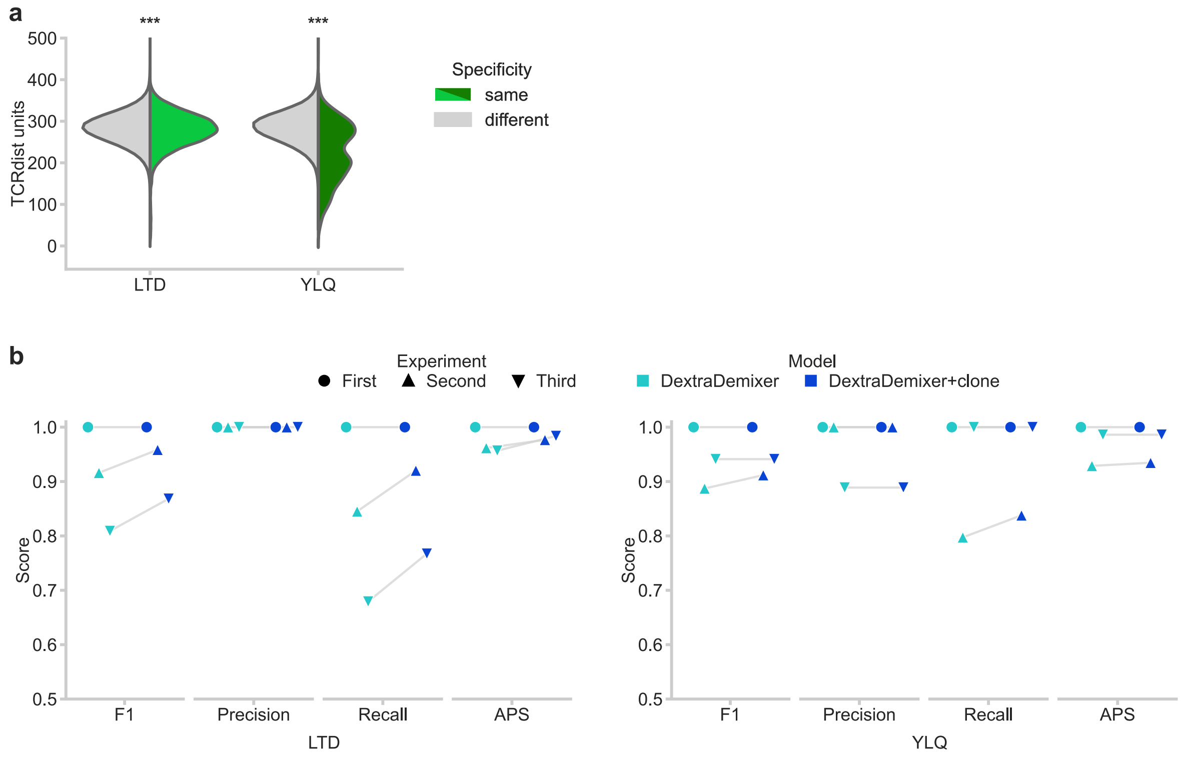


**Supplementary Figure 5 | Additional evaluation of DextraDemixer on SARS-CoV-2 single-cell experiments.** **a,** Distribution of pairwise TCRdist values between clones of the same predicted specificity and between antigen-specific clones to the remaining clones. Statistical significance was tested by an unpaired, two-sided t-test (***: $p<0.001$). **b,** Classification metrics per experiment based on external labels comparing DextraDemixer's base model to the clonotype extension.

### Supplementary Tables

**Supplementary Table 1**: Detailed results on the simulation studies.

**Supplementary Table 2**: Detailed results of all models on the experimental benchmark Gemünd et al.

**Supplementary Table 3**: Detailed results on the longitudinal SARS-CoV-2 study by Kocher et al.
